# Plant structural and nutritional responses to drought differ among common pasture species

**DOI:** 10.1101/2021.10.24.465597

**Authors:** Karen L. M. Catunda, Amber C. Churchill, Sally A. Power, Haiyang Zhang, Kathryn J. Fuller, Ben D. Moore

**Affiliations:** Hawkesbury Institute for the Environment, Western Sydney University, Locked Bag 1797, Penrith, NSW, 2751, Australia; Department of Ecology, Evolution and Behavior, University of Minnesota, 140 Gortner Laboratory, 1479 Gortner Ave., St. Paul, MN, 55108, USA

**Author notes:** Emails: ACC-, SAP-, HZ-, KJF-, BDM.

**Keywords:** grass, leaf:stem ratio, legume, morphology, productivity.

## Abstract

In the face of a changing climate, research indicates that more frequent and severe drought conditions are critical problems that will constrain production of high-quality forage and influence the performance of grazing animals in the future. In addition, the duration of drought and potential trade-offs between plant morphology and nutritional composition may influence plant drought adaptation strategies across pasture species, and the consequences for forage quality are not well understood. Here we present the results of a study investigating the effects of drought on biomass productivity, dead material, leaf:stem biomass allocation and nutritional composition (whole-plant and tissue-specific) across nine diverse pasture species. For this, we conducted a field experiment exposing species to a 6-month period of simulated severe drought (60% rainfall reduction during winter and spring) and samples were collected at multiple harvests. We found that drought had different, harvest-specific effects on plant biomass structure and nutritional composition among pasture species. The severity of drought impacts on productivity, but not on nutritional quality, increased with drought duration. In general, drought strongly reduced productivity, increased the percentage of dead material and had mixed effects (increases, decreases and no effect) on leaf:stem ratio and concentrations of crude protein, non-structural carbohydrates, neutral detergent fibre and lignin. Changes in plant-level nutritional quality were driven by simultaneous changes in both leaf and stem tissues for most, but not all, species. Our findings may be especially helpful for selection of adapted species/cultivars that could minimize potential drought risks on forage, thereby optimising pasture performance under future drought scenarios.

## 1 INTRODUCTION

Grassland ecosystems, which include managed pastures and rangelands, account for approximately 40% of the Earth’s land area and play a key role in food security due to their important service in supplying feeding sources for grazing livestock (ABARES, 2016; Gibson, 2009; Masters et al., 2019; MLA, 2017). However, efficient feeding of livestock is complicated by seasonal and inter-annual changes in plant growth and production (Chapman et al., 2009; Perera et al., 2020). Regional plant productivity is determined by responses to long-term climate patterns, however, local-scale forage availability can be driven by rainfall and associated soil moisture (Brown et al., 2019; McKeon et al., 2009; Murray-Tortarolo and Jaramillo, 2020). As global warming proceeds, many regions of the world will become unable to fulfill requirements for forage quantity and quality, due to more frequent and intense periods of drought (Perera et al., 2020; Rojas-Downing et al., 2017). This will have important implications for the success of the livestock industry and global food production (Dellar et al., 2018; Dumont et al., 2015; Grant et al., 2014).

Drought increases plant water stress and alters plant physiology (Fay, 2009; Heisler-White et al., 2008), which in turn can change plant structural allocation, morphology and nutritional composition. All of these ultimately impact forage nutritional quality and, consequently, animal nutrition and performance (AbdElgawad et al., 2014; Herrero et al., 2015; Howden et al., 2008). Although many studies have addressed the effects of drought on overall biomass production (Churchill et al., 2020; Perera et al., 2019, 2020), a study gap remains in relation to how drought impacts plant structural allocation of biomass and nutritional composition. Furthermore, few studies have imposed longer-term drought under field conditions where plants may have time to acclimatise to stressors. Such acclimation might require trade-offs between structural and nutritional traits, which may not be observed in shorter studies (Deleglise et al., 2015; Grant et al., 2014; McGranahan and Yurkonis, 2018).

Drought can affect forage nutritional quality via changes in structure (e.g. proportional allocation to leaves, stems and flowers), and via changes in the nutritional composition and digestibility of plant parts (particularly leaves and stems), with the magnitude of impacts dependent on plant developmental stage and the severity and duration of drought (Gray and Brady, 2016; IPCC, 2014). Moderate drought stress can delay plant maturation and growth, causing mild or moderate senescence and increases in leaf:stem ratio (Buxton, 1996). However, whole-plant nutritional quality responses to moderate drought conditions are inconsistent across studies; these include no change or reductions in fibre concentration and no change or slight improvements in both crude protein concentration and digestibility of forage species (Deleglise et al., 2015; Dumont et al., 2015; Kuchenmeister et al., 2013; Staniak and Harasim, 2018). This inconsistency may be explained by differences between plant species, the growing stage of the plant when the drought was imposed, drought duration and by differences in the nature of drought treatments. In contrast, studies of prolonged and/or severe drought stress have reported growth inhibition (lower productivity), accelerated maturation, death of plant tissue and decreased leaf:stem ratios (Bruinenberg et al., 2002; Ren et al., 2016). Accompanying these responses are increases in whole-plant fibre concentrations, especially for the lignin fraction, and increased cell-wall thickness and forage toughness, thus reducing forage nutritional quality (Bruinenberg et al., 2002; Deetz et al., 1996; Dumont et al., 2015; Ren et al., 2016). Some studies have also reported reduced quality through decreased concentrations of crude protein and non-structural carbohydrates due to increased translocation of nitrogen and soluble carbohydrates from leaves to roots as senescence proceeds (Buxton, 1996; Durand et al., 2010). While these changes to plant structure and nutritional composition are generally reported separately (Deleglise et al., 2015; Dumont et al., 2015; Ren et al., 2016), herbivores experience their combined consequences. The net impact of these changes on the nutrition of grazers is, however, relatively unknown.

Severe drought generally produces a decrease in forage nutritional quality at the whole-plant level (Buxton, 1996; Deleglise et al., 2015; Durand et al., 2010; Ren et al., 2016), and patterns of resource allocation among plant parts likely underlie many of these changes (Grev et al., 2020). Because some grazers can forage selectively on different plant parts, to various extents, changes to the quality of particular tissues will directly impact herbivores in different ways. There is some evidence that drought differentially affects the nutritional quality of leaves and stems (Pecetti et al., 2017; Wilson et al., 1983). Changes to the relative proportions of plant fractions and to nutritional composition within leaf and stem tissues may reflect diverse adaptation strategies of plants to water stress and strategies to maintain growth (Buxton and Fales, 1994; Le Gall et al., 2015). Understanding these strategies can help to identify plant traits that confer high drought tolerance on plants whilst maintaining structural and nutritional features that ameliorate effects on animal performance under drought conditions (Cavalcante et al., 2014; Tadielo et al., 2017).

This study aimed to investigate the effects of severe drought on pasture productivity, nutritional composition at the whole-plant, leaf and stem levels, the percentage of dead plant material and the leaf:stem biomass ratio. To do this, we conducted a field study exposing nine common pasture species to a 6-month period of severe drought (60% rainfall reduction) during winter and spring, with samples collected across multiple harvests. We hypothesised that drought would reduce forage production and nutritional quality, with drought duration and species-specific differences in the magnitude of effects due to trade-offs in resource allocation among plant parts, such as shifts in leaf:stem biomass ratios.

## 2 MATERIAL AND METHODS

### 2.1 Site description

This study was conducted at the Pastures and Climate Extremes (PACE) facility at the Hawkesbury Campus of Western Sydney University, in Richmond, NSW, Australia (S33.610, E150.740, elevation 25 m; Churchill et al., 2020). The mean annual precipitation at this location is 800 mm (Australian Government Bureau of Meteorology, Richmond - UWS Hawkesbury Station 1980-2010); however, there is large inter-annual variability (between 500 mm and over 1400 mm over the past 30 years). Winter/spring precipitation accounts for 40% of annual rainfall. The mean annual temperature is 17.2 °C, with the warmest and coolest months occurring in January (mean temperature of 22.9 °C) and July (10.2 °C), respectively. The soil is loamy sand with a volumetric water holding capacity of 15-20%, pH of 5.7, plant available nitrogen of 46 mg kg^-1^, plant available (Bray) phosphorus of 26 mg kg^-1^ and 1% soil organic carbon (Churchill et al., 2020). The field facility comprises six replicate polytunnel rainout shelters (48 m x 8 m) with eight treatment plots (4 m x 4 m) per shelter. Individual treatment plots were further subdivided into four subplots, each with a different monoculture or mixed-species sward (total of 192 subplots). This study focuses on all monoculture pasture subplots that were exposed to control and drought treatments, for a total of 108 subplots with nine different pasture species. A detailed overview of the experimental facility descriptions is reported in Churchill et al. (2020).

### 2.2 Selection and establishment of pasture species

Monoculture subplots encompassed a range of functional diversity (C3/C4 grasses, legumes, annuals and perennials) and species’ origins (native grasses, tropical and temperate pastures; **Table 1**) that are all either commonly used in improved grasslands (pastures) or in rangelands across southern Australia and internationally, with the exception of the native grass *Rytidosperma caespitosum*. All pastures were established prior to winter (*Chloris, Digitaria, Festuca* and *Themeda*) or spring (remaining species) of 2018 (Churchill et al., 2020) and swards were managed with seasonal fertilizer application to replace nutrients removed from the soil (55 kg/ha; Cal-Gran Aftergraze, Incitec Pivot Fertilisers, Australia) and hand-weeding to maintain target species dominance. The two legume species received appropriate rhizobium inoculant during sward establishment: ALOSCA granular inoculant for *Biserrula* subplots (Group BS; ALOSCA Technologies, Western Australia, Australia) and EasyRhiz™ soluble legume inoculant and protecting agent for *Medicago* subplots (Group AL; New Edge Microbials, New South Wales, Australia).

**Table 1.**
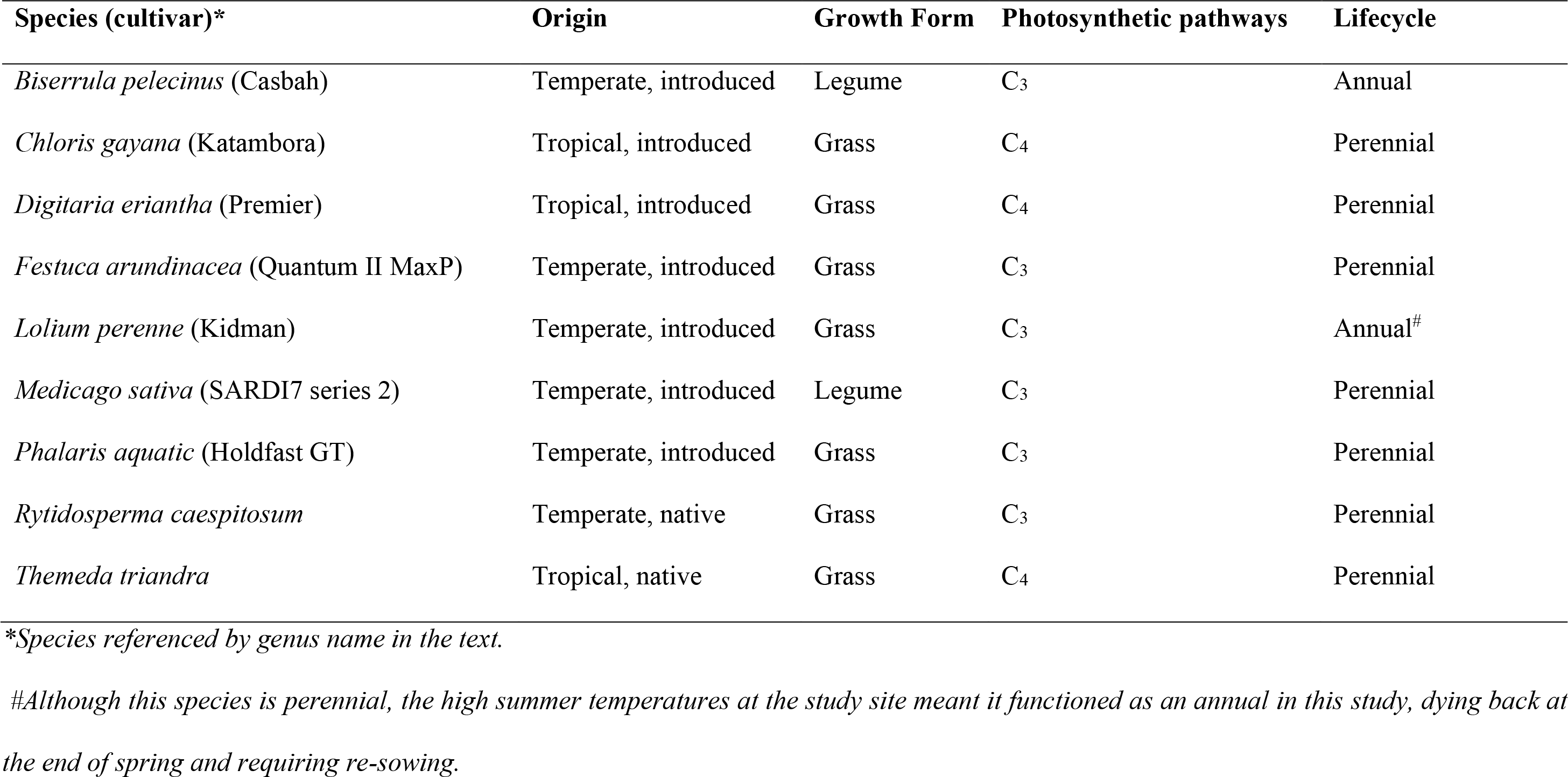
Information about pasture species included in the study.

| Species (cultivar)* | Origin | Growth Form | Photosynthetic pathways | Lifecycle |
| --- | --- | --- | --- | --- |
| *Biserrula pelecinus* (Casbah) | Temperate, introduced | Legume | C<sub>3</sub> | Annual |
| *Chloris gayana* (Katambora) | Tropical, introduced | Grass | C<sub>4</sub> | Perennial |
| *Digitaria eriantha* (Premier) | Tropical, introduced | Grass | C<sub>4</sub> | Perennial |
| *Festuca arundinacea* (Quantum II MaxP) | Temperate, introduced | Grass | C<sub>3</sub> | Perennial |
| *Lolium perenne* (Kidman) | Temperate, introduced | Grass | C<sub>3</sub> | Annual<sup>#</sup> |
| *Medicago sativa* (SARDI7 series 2) | Temperate, introduced | Legume | C<sub>3</sub> | Perennial |
| *Phalaris aquatic* (Holdfast GT) | Temperate, introduced | Grass | C<sub>3</sub> | Perennial |
| *Rytidosperma caespitosum* | Temperate, native | Grass | C<sub>3</sub> | Perennial |
| *Themeda triandra* | Tropical, native | Grass | C<sub>4</sub> | Perennial |

\*Species referenced by genus name in the text.
#Although this species is perennial, the high summer temperatures at the study site meant it functioned as an annual in this study, dying back at the end of spring and requiring re-sowing.

### 2.3 Experimental treatments and environmental monitoring

All nine pasture species were exposed to the same irrigation regime. The control (C) treatment represented a typical precipitation regime for the local area during years with annual precipitation between 650-750 mm, accounting for long-term patterns in seasonality and in the statistical distribution of event sizes and timing within seasons. In the drought (D) treatment, precipitation event sizes were reduced by 60% throughout the 6-month austral winter/spring period from 1 June to 30 November 2019. This drought treatment represented the drier end of climate model predictions for end-of-century seasonal rainfall change for southeastern Australia, under the Representative Concentration Pathway - RCP8.5 (CSIRO, 2020). A 60% reduction in rainfall falls within the range of observed historical rainfall patterns for key pasture growing regions across southeastern Australia, including the study site, and such extremes are predicted to increase in frequency and duration (BOM, 2019). Target precipitation was applied using an irrigation system installed in each plot (5 irrigation points in each) as described in Churchill et al. (2020). Prior to the start of the winter season, all plots received the same irrigation inputs (1 December 2018 to 31 May 2019; 419.7 mm total amount).

Environmental monitoring of treatment plots included continuous recording of soil moisture (0-15 cm; 16 per shelter; Time Domain Reflectometers; CS616, Campbell Scientific) in four different species subplots (*Biserrula*, *Festuca*, *Lolium, Medicago*). Air temperature and humidity sensors (Series RHP-2O3B, Dwyer Instruments Inc, USA) mounted in force- ventilated radiation shields were installed inside and outside the rainout shelters at 60 cm height, with records collected every 5 min to determine shelter effects on environmental conditions. The amount of irrigation applied in each treatment, air temperature and soil moisture averaged across the shelters during the 6-month experimental period (1 June to 30 November 2019) can be seen in **Supplementary Figure S1**.

### 2.4 Plant sampling during the experimental period and measurements

All subplots were managed and harvested regularly before and during this study based on grazing system recommendations practiced in the study region (Clements et al., 2003). Harvesting involved the use of hand shears and a sickle mower. Prior to the start of the winter season, all species were harvested at the end of May 2019. During the 6-month winter/spring experimental drought period, aboveground productivity was determined via three harvests, one in mid-August, one in early October and one in mid-November 2019, for all species, except for *Chloris* and *Digitaria* that in the August harvest there was no plant biomass in the plots of these species (**Supplementary Table S1**). In all harvests, plants were cut to 5 cm above the soil surface and weighed (fresh mass), with a representative sub-sample sorted to remove/exclude weeds and to determine the percentage of dead material in the total biomass by weight (fresh mass); thereafter, all plant biomass sub-samples, including live and dead material, were immediately microwaved at 600W for 90 seconds to stop enzymatic activity (Landhäusser et al., 2018) and then oven-dried at 65 °C for at least 48 hours (until constant weight), and weighed to determine total dry matter productivity (kg DM ha^-1^; live and dead material) per harvest, for each species and treatment.

### 2.5 Plant structural analysis and sample processing

For the nutritional analysis of the whole-plant material, we analysed dry samples from the August, October and November harvests, which were composed of a proportionally representative mixture of live and dead leaves, stems (or culms/tillers) and inflorescences (**Supplementary Table S1**). In addition, for the November harvest, we sorted samples (composed of both live and dead material) into leaves and stems (or culms/tillers), weighed these fractions to calculate the leaf:stem ratio (**Supplementary Table S1**) and analysed the nutritional composition of the fractions separately. Dried samples were ground through a 1-mm screen in a laboratory mill (Foss Cyclotec Mill, Denmark) and stored in airtight plastic containers in the dark at room temperature prior to collection of near-infrared reflectance (NIR) spectra and wet chemical analysis. Plant samples were further homogenized using a ball-mill to produce a fine powder prior to nitrogen determination by elemental analysis (Retsch® MM200; Hann, Germany).

### 2.6 Nutritional analysis

All dried and ground samples were scanned twice and their spectra were averaged using a near-infrared reflectance spectrophotometer (NIRS; FOSS XDS Rapid Content™ Analyzer) with a spectral range of 400 to 2500 nm. Representative samples were selected using the software WinISI 4.8.0 (FOSS Analytical A/S, Denmark) for analysis of nutrient composition by wet chemistry for all parameters, in order to calibrate and validate the NIR.

The selected samples were analysed for ash (ASH) according to the standard methods and procedures for animal feed outlined by the Association of Official Analytical Chemists (AOAC, 1990). Nitrogen (N) concentration was determined from ∼ 100 mg samples using an automated combustion method on a Leco TruMac CN analyzer (Leco Corporation, USA). Crude protein (CP) concentration was then calculated by applying a 6.25 conversion factor to the N concentration (AOAC, 1990). Ether extract (EE) was determined according to the American Oil Chemists’ Society-AOCS high-temperature method using petroleum ether (B.P. 40-70 °C) and the Soxhlet method (Buchi 810 Soxhlet Multihead Extract Rack, UK). Fibre fractions were determined with an ANKOM Fibre Analyzer (model 200, ANKOM® Technology, NY, USA) with the use of neutral and acid detergent solutions and corrected for dry matter content (Goering and Van Soest, 1970). Samples were analysed for neutral detergent fibre (NDF), acid detergent fibre and acid detergent lignin (ADL) by the sequential method of Van Soest and Robertson (1980). Sodium sulphite and α-amylase were added to the solution for NDF determination. The values of ASH, EE, CP and NDF were used to calculate non-structural carbohydrates (NSC) according to Sniffen et al. (1992). All nutritional parameters were expressed as a percentage of total DM.

Details associated with mathematical treatment of spectra and descriptive statistics for NIRS calibration can be found in Catunda et al. (2021). However, in brief, for the development of NIRS calibration models, modified Partial Least Squares regression with cross-validation was used to develop predictive equations for each nutritional parameter to prevent overfitting of models (Shenk and Westerhaus 1991; Catunda et al., 2021). The NIRS calibration equations were considered to be both suitable and robust to estimate all the nutritional parameters of the samples of all pasture species assessed (Catunda et al., 2021).

### 2.7 Calculations and statistical analysis

We analysed the effects of drought on pasture productivity, percentage of dead material and nutritional composition of the whole-plant separately for each harvest, but only considered changes in leaf:stem ratio and nutritional composition of leaf and stem fractions at the end of the drought period (November). All pasture responses were analysed using linear mixed-effects (LME) models in the ‘lme4’ package in the software R version 4.0.0 (R Core Team, 2020; Bates et al., 2015). Watering regime (Control: C, Drought: D) was included as a fixed effect and the rainout shelter as a random factor; residuals were checked for normality. We calculated the mean effect size due to drought (*Equation 1*; for the figures in the results section) as the ratio of drought to their respective control treatment values, along with 95% confidence intervals (CI).

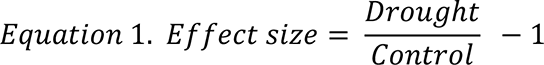

In the effect size figures, positive values represent responses that are greater under drought than in control plots, while negative values represent the opposite. We expressed effect sizes as percentages (effect size multiplied by 100) in the text throughout the results section.

Finally, to produce a more holistic overview of effects of drought on plant response variables across all pasture species, we performed a principal component analysis (PCA) of the data from the end of the drought period (November harvest). To test for the effects of the watering regime on plant responses, we undertook permutational analysis of variance (PERMANOVA) using the ‘vegan’ package (Oksanen et al., 2020) in R version 4.0.0 (R Core Team, 2020).

## 3 RESULTS

### 3.1 Effects of drought on productivity, dead material and nutritional composition of the whole-plant

The effect of drought on dry matter productivity, percentage of dead material and nutritional composition varied among the nine pasture species studied and, in some cases, also differed between individual harvests. These effects included either a significant reduction or no effect on productivity and an increase or no effect on the percentage of dead material. Drought had varied effects on the whole-plant concentrations of CP, NSC, NDF and ADL for different species and harvests (**Table 2**; **Figure 1**).

**Figure 1.**
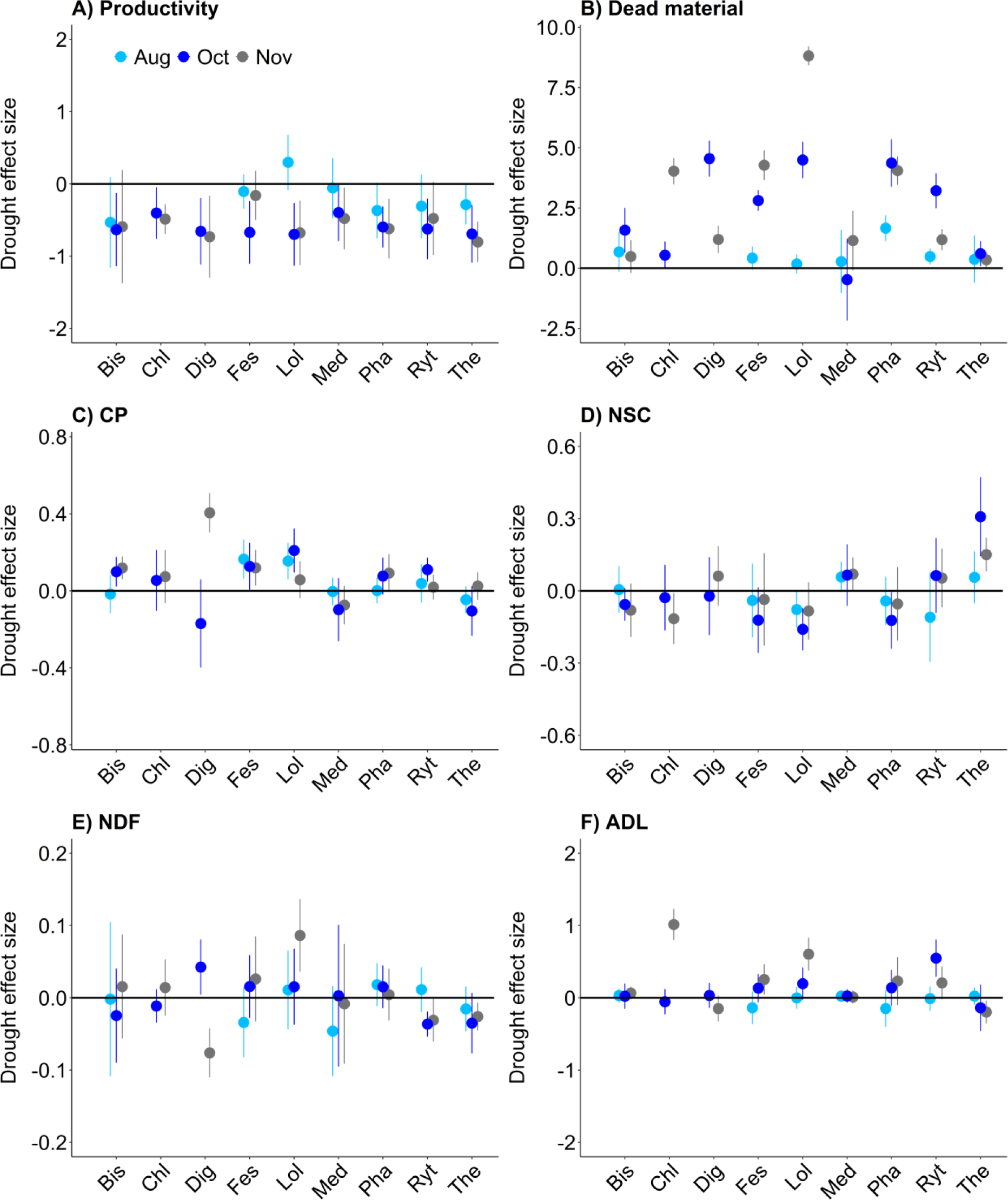
Drought effect sizes on A) productivity, B) percentage of dead material, and whole-plant nutritional composition [C) crude protein, D) non-structural carbohydrates, E) neutral detergent fibre, F) acid detergent lignin] of pasture species during the drought period (August, October and November harvests separately). Values shown are means with vertical bars representing 95% confidence intervals (n = 6). Species abbreviations are as follows: *Biserrula* (Bis), *Chloris* (Chl), *Digitaria* (Dig), *Festuca* (Fes), *Lolium* (Lol), *Medicago* (Med), *Phalaris* (Pha), *Rytidosperma* (Ryt), *Themeda* (The). Note: during the harvest in August, there was no biomass for *Chloris* and *Digitaria*.

**Table 2.** Mean ± standard errors (n = 6) for productivity (kg DM ha^-1^), percentage of dead material, and nutritional composition (in percentage of dry matter) of the whole-plant of pasture species grown under different watering regimes (control, C; drought, D) treatments during the drought period (August, October and November harvests).

| Species | Harvest | Productivity |  | Dead material |  | CP |  | NSC |  | NDF |  | ADL |  |
| --- | --- | --- | --- | --- | --- | --- | --- | --- | --- | --- | --- | --- | --- |
|  |  | C | D | C | D | C | D | C | D | C | D | C | D |
| *Bis* | August | 1737 ± 376 | **812 ± 189\*** | 2.2 ± 0.7 | 3.6 ± 0.9 | 19.2 ± 0.6 | 18.9 ± 0.8 | 29.3 ± 0.6 | 29.5 ± 1.3 | 38.3 ± 0.7 | 38.2 ± 2.0 | 7.9 ± 0.2 | 8.1 ± 0.3 |
| | October | 882 ± 177 | **323 ± 52.1\*** | 1.7 ± 0.7 | **4.3 ± 0.9\*** | 16.1 ± 0.3 | **17.7 ± 0.6\*** | 30.9 ± 0.9 | 29.1 ± 0.4 | 42.2 ± 1.1 | 41.2 ± 0.8 | 7.6 ± 0.5 | 7.7 ± 0.4 |
| | November | 304 ± 83.8 | 124 ± 35.7 | 6.9 ± 1.1 | 10.3 ± 3.1 | 14.6 ± 0.3 | **16.3 ± 0.3\*** | 30.8 ± 0.7 | 28.3 ± 1.5 | 44.7 ± 0.9 | 45.4 ± 1.4 | 7.6 ± 0.2 | 8.1 ± 0.3 |
| *Chl* | October | 1176 ± 137 | 702 ± 96.5 | 7.2 ± 1.1 | 11.2 ± 2.7 | 6.3 ± 0.4 | 6.6 ± 0.3 | 17.5 ± 0.9 | 17.0 ± 0.7 | 67.3 ± 0.6 | 66.5 ± 0.5 | 4.3 ± 0.2 | 4.0 ± 0.3 |
| | November | 1646 ± 168 | **844 ± 12.7\*** | 6.2 ± 1.6 | **31.3 ± 3.1\*** | 6.7 ± 0.3 | 7.2 ± 0.4 | 17.3 ± 0.7 | 15.3 ± 0.5 | 65.6 ± 0.7 | 66.5 ± 1.1 | 2.9 ± 0.1 | **5.9 ± 0.6\*** |
| *Dig* | October | 1128 ± 109 | **389 ± 82.5\*** | 9.6 ± 1.9 | **53.3 ± 17\*** | 11.6 ± 0.8 | **9.6 ± 0.9\*** | 12.9 ± 0.9 | 12.6 ± 0.5 | 62.6 ± 0.8 | **65.2 ± 0.9\*** | 6.2 ± 0.3 | 6.4 ± 0.4 |
| | November | 2535 ± 217 | **681 ± 187\*** | 6.7 ± 1.5 | **14.6 ± 2.7\*** | 7.1 ± 0.3 | **10.0 ± 0.2\*** | 15.0 ± 0.8 | 15.9 ± 0.5 | 67.1 ± 0.8 | **62.0 ± 0.8\*** | 3.8 ± 0.3 | 3.3 ± 0.2 |
| *Fes* | August | 1204 ± 133 | 1076 ± 52.5 | 7.1 ± 0.4 | 10.1 ± 2.4 | 12.1 ± 0.5 | **14.0 ± 0.4\*** | 21.7 ± 1.1 | 20.8 ± 1.2 | 54.2 ± 0.5 | 52.4 ± 1.2 | 4.5 ± 0.1 | 3.9 ± 0.4 |
| | October | 1001 ± 117 | **328 ± 61.1\*** | 6.5 ± 1.1 | **24.6 ± 3.1\*** | 11.6 ± 0.5 | 13.0 ± 0.5 | 19.7 ± 0.5 | 17.3 ± 1.1 | 57.1 ± 0.8 | 58.0 ± 1.0 | 4.3 ± 0.3 | 4.9 ± 0.3 |
| | November | 612 ± 68.3 | 513 ± 67.6 | 12.6 ± 3.9 | **66.6 ± 3.3\*** | 11.0 ± 0.4 | **12.3 ± 0.3\*** | 17.3 ± 1.4 | 16.7 ± 0.9 | 56.7 ± 1.5 | 58.2 ± 0.8 | 6.0 ± 0.6 | **7.5 ± 0.4\*** |
| *Lol* | August | 2085 ± 328 | 2707 ± 311 | 2.5 ± 0.3 | 3.0 ± 0.5 | 8.4 ± 0.3 | **9.7 ± 0.3\*** | 39.1 ± 0.5 | 36.1 ± 1.3 | 41.9 ± 0.7 | 42.4 ± 0.9 | 2.8 ± 0.1 | 2.8 ± 0.2 |
| | October | 1110 ± 163 | **335 ± 55.0\*** | 3.0 ± 0.9 | **16.6 ± 4.0\*** | 11.8 ± 0.6 | **14.3 ± 0.4\*** | 25.2 ± 0.3 | **21.2 ± 0.9\*** | 51.2 ± 0.8 | 52.0 ± 1.1 | 3.5 ± 0.3 | 4.2 ± 0.3 |
| | November | 599 ± 108 | **192 ± 26.3\*** | 5.1 ± 0.8 | **50.3 ± 6.3\*** | 11.2 ± 0.4 | 11.9 ± 0.4 | 20.9 ± 0.9 | 19.1 ± 0.8 | 53.7 ± 1.0 | **58.3 ± 1.0\*** | 3.2 ± 0.3 | **5.2 ± 0.4\*** |
| *Med* | August | 1503 ± 230 | 1417 ± 204 | 1.6 ± 0.6 | 2.1 ± 1.1 | 15.3 ± 0.4 | 15.2 ± 0.4 | 30.7 ± 0.6 | 32.5 ± 0.8 | 42.6 ± 0.9 | 40.6 ± 0.9 | 9.4 ± 0.3 | 9.6 ± 0.3 |
| | October | 2610 ± 275 | **1577 ± 271\*** | 1.0 ± 0.4 | 1.0 ± 0.1 | 17.2 ± 0.9 | 15.5 ± 1.0 | 27.4 ± 1.5 | 29.2 ± 1.1 | 44.3 ± 1.1 | 44.5 ± 1.9 | 8.8 ± 0.2 | 9.0 ± 0.4 |
|  | November | 2732 ± 400 | **1425 ± 226\*** | 1.0 ± 0.2 | **1.3 ± 0.7\*** | 15.5 ± 0.7 | 14.4 ± 0.3 | 28.7 ± 0.9 | 30.7 ± 0.6 | 44.7 ± 1.7 | 44.4 ± 0.8 | 8.5 ± 0.2 | 8.5 ± 0.3 |
| *Pha* | August | 1817 ± 284 | **1148 ± 135\*** | 1.7 ± 0.4 | **4.5 ± 0.8\*** | 14.1 ± 0.4 | 14.1 ± 0.3 | 26.5 ± 1.2 | 25.4 ± 0.6 | 45.9 ± 0.5 | 46.8 ± 0.5 | 2.9 ± 0.2 | 2.4 ± 0.3 |
|  | October | 1299 ± 130 | **523 ± 54.2\*** | 1.4 ± 0.4 | **7.4 ± 3.0\*** | 15.3 ± 0.7 | 16.5 ± 0.3 | 18.2 ± 0.9 | 16.0 ± 0.5 | 53.1 ± 0.8 | 53.9 ± 0.3 | 2.9 ± 0.3 | 3.3 ± 0.1 |
|  | November | 621 ± 72.8 | **236 ± 41.0\*** | 5.9 ± 1.5 | **29.7 ± 4.5\*** | 9.7 ± 0.4 | 10.6 ± 0.3 | 16.2 ± 0.4 | 15.3 ± 1.1 | 61.9 ± 0.9 | 62.2 ± 0.7 | 4.0 ± 0.3 | 4.9 ± 0.7 |
| *Ryt* | August | 2081 ± 397 | 1439 ± 169 | 2.8 ± 0.4 | 4.2 ± 0.3 | 11.0 ± 0.5 | 11.5 ± 0.3 | 14.7 ± 1.0 | 13.1 ± 0.9 | 63.5 ± 0.8 | 64.2 ± 0.5 | 4.0 ± 0.3 | 3.9 ± 0.2 |
|  | October | 1932 ± 317 | **727 ± 98.4\*** | 2.1 ± 0.7 | **9.0 ± 1.6\*** | 10.8 ± 0.2 | 12.0 ± 0.3 | 10.1 ± 0.4 | 10.8 ± 0.8 | 69.8 ± 0.3 | **67.3 ± 0.5\*** | 2.2 ± 0.2 | **3.4 ± 0.3\*** |
|  | November | 618 ± 123 | 323 ± 52.8 | 31.0 ± 5.7 | **67.7 ± 8.4\*** | 9.9 ± 0.3 | 10.1 ± 0.2 | 13.2 ± 0.6 | 13.9 ± 0.6 | 68.4 ± 0.9 | 66.3 ± 0.5 | 5.3 ± 0.6 | **6.4 ± 0.3\*** |
| *The* | August | 2764 ± 241 | 1972 ± 213 | 2.9 ± 0.8 | 4.0 ± 1.7 | 9.5 ± 0.3 | 9.1 ± 0.2 | 16.0 ± 0.7 | 16.9 ± 0.5 | 65.8 ± 0.9 | 64.8 ± 0.5 | 4.9 ± 0.2 | 5.0 ± 0.2 |
|  | October | 3685 ± 272 | **1136 ± 214\*** | 1.0 ± 0.1 | 1.0 ± 0.1 | 9.1 ± 0.4 | 8.1 ± 0.3 | 13.3 ± 0.8 | **17.4 ± 1.0\*** | 70.2 ± 0.7 | 67.7 ± 1.3 | 3.7 ± 0.3 | 3.2 ± 0.5 |
|  | November | 1917 ± 130 | **379 ± 46.9\*** | 3.3 ± 0.4 | 4.5 ± 0.4 | 7.2 ± 0.1 | 7.4 ± 0.2 | 15.8 ± 0.4 | **18.2 ± 0.5\*** | 69.7 ± 0.5 | 67.9 ± 0.4 | 4.4 ± 0.1 | **3.5 ± 0.3\*** |

Note: Asterisks (\*) and **bold** values denote statistical significance at the $p \leq 0.05$ level. During the harvest in August, there was no biomass for *Chloris* and *Digitaria* under both treatments.
Abbreviations: CP: crude protein; NSC: non-structural carbohydrates; NDF: neutral detergent fibre; ADL: acid detergent lignin; Bis: *Biserrula*; Chl: *Chloris*; Dig: *Digitaria*; Fes: *Festuca*; Lol: *Lolium*; Med: *Medicago*; Pha: *Phalaris*; Ryt: *Rytidosperma*; The: *Themeda*.

Across species and harvests, productivity ranged from 304 (*Biserrula*, November) to 3,685 kg DM ha^-1^ (*Themeda*, October) under control treatment and the peak productivity varied across species, for example, *Themeda* peaked in October and *Biserrula* in August (**Table 2**). Total productivity under the control treatment across the six-month period was greatest for *Themeda* (8,366 kg DM ha^-1^) and *Medicago* (6,845 kg DM ha^-1^) and lowest for *Chloris* (2,822 kg DM ha^-1^) and *Festuca* (2,817 kg DM ha^-1^). Droughted subplots were significantly less productive, and drought impacts became progressively greater across harvests, with the last two harvests being the most affected (**Figure 1A, Supplementary Table S2**). All pasture species were strongly and significantly affected by the drought, but not all species were significantly affected at all harvests. The greatest reductions in productivity per harvest were seen for *Themeda* (-80% November) and *Digitaria* (-73% November), and the smallest for *Medicago* and *Chloris* (both -48%, November; **Figure 1A**).

The mean percentage of dead material ranged from 1% to 31% in control subplots, and from 1% to 68% in droughted plots (**Table 2**). In contrast to productivity responses, the severity of drought impacts on the percentage of dead material was not generally progressive across harvests, and for some species, in fact, was actually reduced in successive harvests. Overall, the percentage of dead material increased under drought for all species, except *Themeda*, which was not affected in any harvest (*p* > 0.05; **Figure 1B; Supplementary Table S2**). The most strongly affected species were *Lolium* (+886%, November), *Phalaris* (+429%, October) and *Festuca* (+429%, November). And the least affected species were *Biserrula* (+153%, October) and *Medicago* (+117%, November; **Figure 1B**).

Effects of drought on whole-plant nutritional composition were apparent for seven of the nine study species. The exceptions were *Medicago* and *Phalaris*, which, in fact, experienced no significant treatment impacts on nutritional quality at any time throughout the experiment (**Table 2; Supplementary Table S2**). Drought effects on nutritional parameters are summarised in **Figure 1C-F**. Overall, the severity of drought impacts on forage nutritional quality was not generally progressive across harvests. In the August harvest, drought only had impacts on *Festuca* and *Lolium*, which experienced an increase in CP (+15% for both species). In October, the drought increased CP (+10%) in *Biserrula* and NSC (+31%) in *Themeda*, but decreased CP (-17%) and slightly increased NDF (+4%) in *Digitaria*. In November, drought was associated with improved nutritional quality in *Digitaria* (+41% CP and -8% NDF) and *Themeda* (+15% NSC and -20% ADL), but reduced the nutritional quality of *Lolium* through an increase in NDF (+9%) and ADL (+63%).

### 3.2 Effects of drought on leaf:stem ratio and nutritional composition of leaf and stem tissues

At the end of the 6-month period of drought (November harvest), the drought treatment significantly increased the leaf:stem ratio of *Phalaris* by 129%, *Themeda* by 102%, and *Digitaria* by 80%, and decreased that of *Chloris* by 50% (**Table 3**; **Figure 2A**). However, drought had no effect on the leaf:stem ratios of the remaining species (*p* > 0.05; **Supplementary Table S3**).

**Figure 2.**
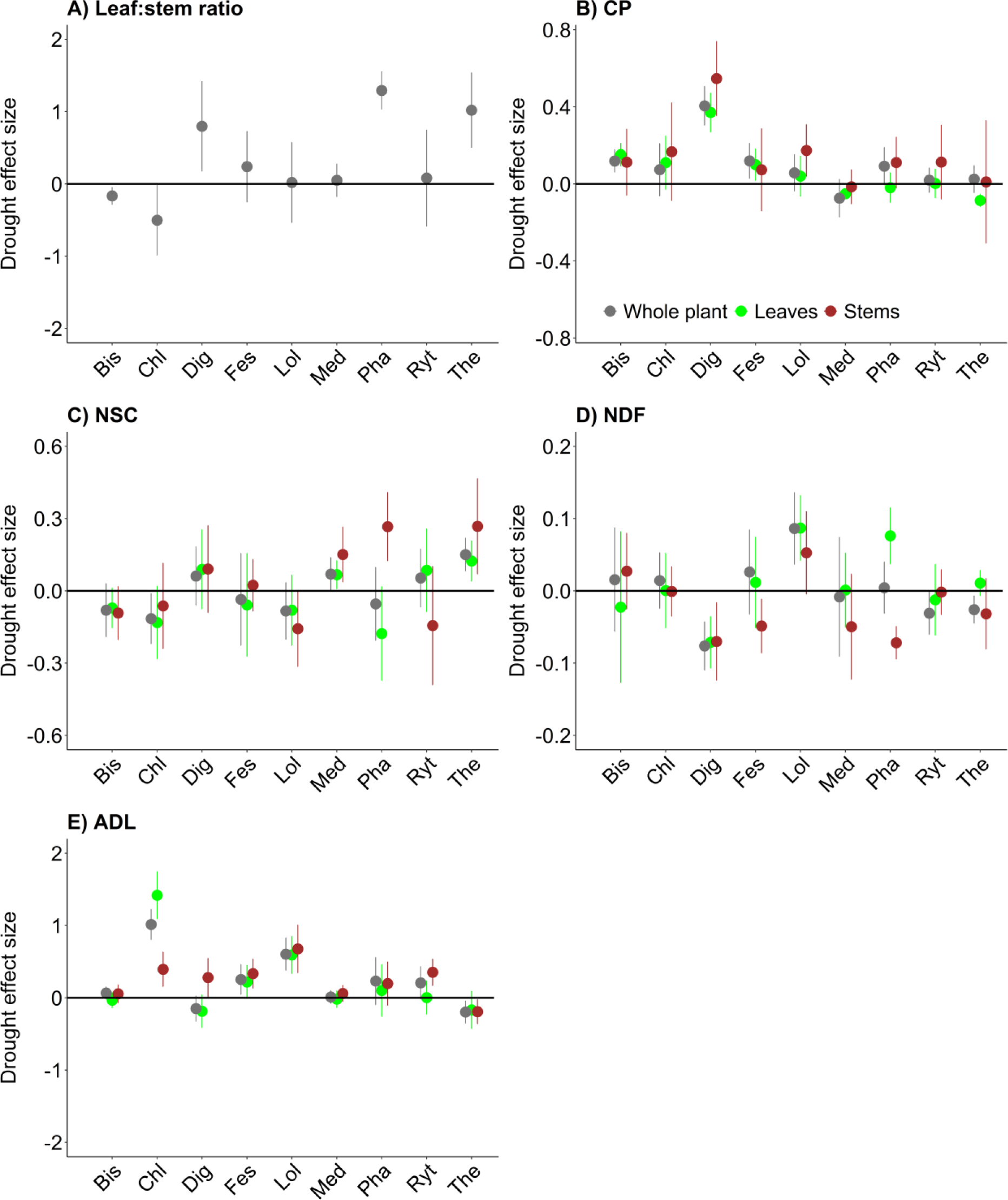
Drought effect sizes on leaf:stem ratio and nutritional composition [A) crude protein, B) non-structural carbohydrates, C) neutral detergent fibre, D) acid detergent lignin] of whole- plant (grey circle), leaves (green circle) and stems (brown circle) of pasture species at the end of the drought period (November harvest). Values shown are means with vertical bars representing 95% confidence intervals (n = 6). Species abbreviations follow Figure 1.

**Table 3.** Mean ± standard errors (n = 6) for leaf:stem ratio and nutritional composition of plant parts (leaves and stems) of pasture species grown under different watering regimes (control, C; drought, D) treatments at the end of the drought period (November harvest).

| Species | Leaf:stem |  | Plant parts | Parameters (in percentage of dry matter) |  |  |  |  |  |  |  |
| --- | --- | --- | --- | --- | --- | --- | --- | --- | --- | --- | --- |
|  |  |  |  | CP |  | NSC |  | NDF |  | ADL |  |
|  | C | D |  | C | D | C | D | C | D | C | D |
| Bis | 1.6 ± 0.1 | 1.3 ± 0.1 | Leaves | 17.1 ± 0.4 | 19.7 ± 0.3* | 38.1 ± 0.5 | 35.4 ± 1.4 | 32.7 ± 1.0 | 31.9 ± 1.4 | 8.1 ± 0.3 | 7.8 ± 0.3 |
|  |  |  | Stems | 10.6 ± 0.7 | 11.8 ± 0.7 | 28.7 ± 1.3 | 26.1 ± 0.9 | 51.5 ± 1.2 | 52.9 ± 0.7 | 9.4 ± 0.4 | 9.9 ± 0.5 |
| Chl | 4.1 ± 0.9 | 2.0 ± 0.2* | Leaves | 7.3 ± 0.4 | 8.1 ± 0.4 | 17.1 ± 0.7 | 14.9 ± 1.0 | 64.1 ± 0.7 | 64.2 ± 1.5 | 2.5 ± 0.2 | 6.0 ± 0.9* |
|  |  |  | Stems | 4.7 ± 0.5 | 5.5 ± 0.5 | 17.7 ± 1.3 | 16.5 ± 0.9 | 70.9 ± 1.1 | 70.8 ± 0.6 | 4.2 ± 0.4 | 5.9 ± 0.4* |
| Dig | 4.8 ± 1.1 | 8.7 ± 1.8* | Leaves | 7.6 ± 0.4 | 10.4 ± 0.2* | 14.7 ± 1.1 | 16.1 ± 0.7 | 65.5 ± 0.9 | 60.9 ± 0.7* | 3.9 ± 0.3 | 3.2 ± 0.2 |
|  |  |  | Stems | 4.9 ± 0.3 | 7.6 ± 0.6* | 14.5 ± 0.9 | 15.8 ± 1.1 | 74.2 ± 1.2 | 69.0 ± 1.5* | 3.4 ± 0.2 | 4.4 ± 0.5 |
| Fes | 7.7 ± 1.3 | 9.5 ± 1.7 | Leaves | 11.5 ± 0.4 | 12.6 ± 0.3* | 17.2 ± 1.5 | 16.2 ± 1.0 | 57.4 ± 1.6 | 58.1 ± 0.9 | 6.3 ± 0.7 | 7.7 ± 0.4* |
|  |  |  | Stems | 8.9 ± 0.9 | 9.5 ± 0.5 | 20.3 ± 0.9 | 20.8 ± 0.7 | 61.7 ± 0.9 | 58.7 ± 0.7* | 4.8 ± 0.3 | 6.4 ± 0.5* |
| Lol | 7.9 ± 0.5 | 8.1 ± 2.2 | Leaves | 11.8 ± 0.5 | 12.2 ± 0.4 | 20.0 ± 1.0 | 18.4 ± 1.0 | 53.9 ± 0.7 | 58.5 ± 1.1* | 3.3 ± 0.3 | 5.3 ± 0.5* |
|  |  |  | Stems | 8.2 ± 0.5 | 9.6 ± 0.2* | 28.8 ± 1.8 | 24.3 ± 1.3* | 54.5 ± 1.1 | 57.4 ± 1.2 | 2.5 ± 0.4 | 4.3 ± 0.3* |
| Med | 0.7 ± 0.1 | 0.8 ± 0.1 | Leaves | 22.3 ± 0.1 | 21.1 ± 0.2 | 33.7 ± 0.6 | 36.0 ± 0.9 | 30.7 ± 0.5 | 30.7 ± 0.6 | 9.0 ± 0.2 | 8.8 ± 0.5 |
|  |  |  | Stems | 8.9 ± 0.3 | 8.8 ± 0.3 | 22.9 ± 0.9 | 26.4 ± 1.1* | 60.1 ± 1.7 | 57.2 ± 1.4 | 8.1 ± 0.3 | 8.6 ± 0.4 |
| *Pha* | 1.4 ± 0.1 | **3.1 ± 0.4\*** | Leaves | 12.2 ± 0.3 | 12.0 ± 0.4 | 16.7 ± 0.5 | **13.7 ± 1.3\*** | 56.4 ± 0.5 | **60.7 ± 1.1\*** | 4.8 ± 0.4 | 5.3 ± 0.9 |
|  |  |  | Stems | 5.8 ± 0.2 | 6.5 ± 0.4 | 16.0 ± 0.8 | **20.2 ± 1.1\*** | 71.1 ± 0.5 | **66.0 ± 0.6\*** | 3.1 ± 0.4 | 3.7 ± 0.3 |
| *Ryt* | 4.1 ± 1.3 | 4.4 ± 0.6 | Leaves | 11.0 ± 0.3 | 11.0 ± 0.3 | 13.3 ± 1.0 | 14.5 ± 0.6 | 65.5 ± 1.4 | 64.6 ± 0.8 | 6.5 ± 0.7 | 6.5 ± 0.3 |
|  |  |  | Stems | 5.9 ± 0.5 | 6.5 ± 0.4 | 15.0 ± 1.3 | 12.9 ± 1.2 | 73.3 ± 0.9 | 73.1 ± 0.8 | 4.9 ± 0.3 | **6.6 ± 0.4\*** |
| *The* | 1.7 ± 0.3 | **3.4 ± 0.6\*** | Leaves | 9.6 ± 0.1 | 8.8 ± 0.1 | 16.4 ± 0.5 | **18.4 ± 0.5\*** | 64.5 ± 0.4 | 65.2 ± 0.4 | 4.1 ± 0.2 | 3.4 ± 0.4 |
|  |  |  | Stems | 3.3 ± 0.3 | 3.4 ± 0.4 | 11.8 ± 1.0 | **15.0 ± 0.9\*** | 80.5 ± 1.5 | 78.0 ± 1.3 | 4.9 ± 0.4 | **4.0 ± 0.1\*** |

*Note: Asterisks (\*) and **bold** values denote statistical significance at the $p \leq 0.05$ level.*
*Abbreviations: CP: crude protein; NSC: non-structural carbohydrates; NDF: neutral detergent fibre; ADL: acid detergent lignin; Bis: Biserrula;*
*Chl: Chloris; Dig: Digitaria; Fes: Festuca; Lol: Lolium; Med: Medicago; Pha: Phalaris; Ryt: Rytidosperma; The: Themeda.*

Tissue-specific responses to drought varied in both magnitude and direction across pasture species (**Table 3**; **Supplementary Table S3**). For instance, drought increased CP in *Biserrula* and *Festuca* leaf tissue, in both leaves and stems in *Digitaria* and in the stems only in *Lolium* (**Figure 2B**). In contrast, drought decreased NSC in *Lolium* stems while increasing it in both plant parts in *Themeda* (**Figure 2C**). Drought also decreased NDF in *Digitaria* stems and leaves, while increasing it in *Lolium* leaf tissue (**Figure 2D**). In both plant parts of *Chloris*, *Festuca* and *Lolium*, as well as in *Rytidosperma* stems, the drought increased ADL, while in *Themeda* stems it decreased. (**Figure 2E**). Interestingly, *Medicago* and *Phalaris* were the only species where drought affected the nutritional composition of individual plant parts but not the whole-plant. Specifically, in *Medicago,* drought increased NSC in the stem tissue, with no other changes detected. For *Phalaris,* drought affected plant parts in opposite directions for NSC (**Figure 2C**) and NDF (**Figure 2D**).

### 3.4 Assessing plant responses to drought in a multivariate context

The first two principal components explained 73% of the variation in plant responses across treatments (**Figure 3**). We found that drought had a significant effect (PERMANOVA: *p* < 0.01) across all pasture species (**Figure 3A**). Differences in multivariate plant responses among individual species were further apparent with clear separation between responses of legumes (*Biserrula* and *Medicago)* and grasses (PC1; **Figure 3B**), and within grasses, with C3 grasses differing from C4 grasses along PC2. The first principal component (PC1, 47.9% data variance) was associated with nutritional composition and had positive loadings for CP, ADL and NSC, and negative loadings for NDF (**Figure 3C**). The second component (PC2, 24.7% data variance) was associated with plant structural characteristics, including positive loadings for the percentage of dead material and leaf:stem ratio, and negative loadings for total biomass production. Overall, nutritional parameters explained a greater proportion of the variance of all measured responses than morphological parameters across treatments and all studied pasture species.

**Figure 3.**
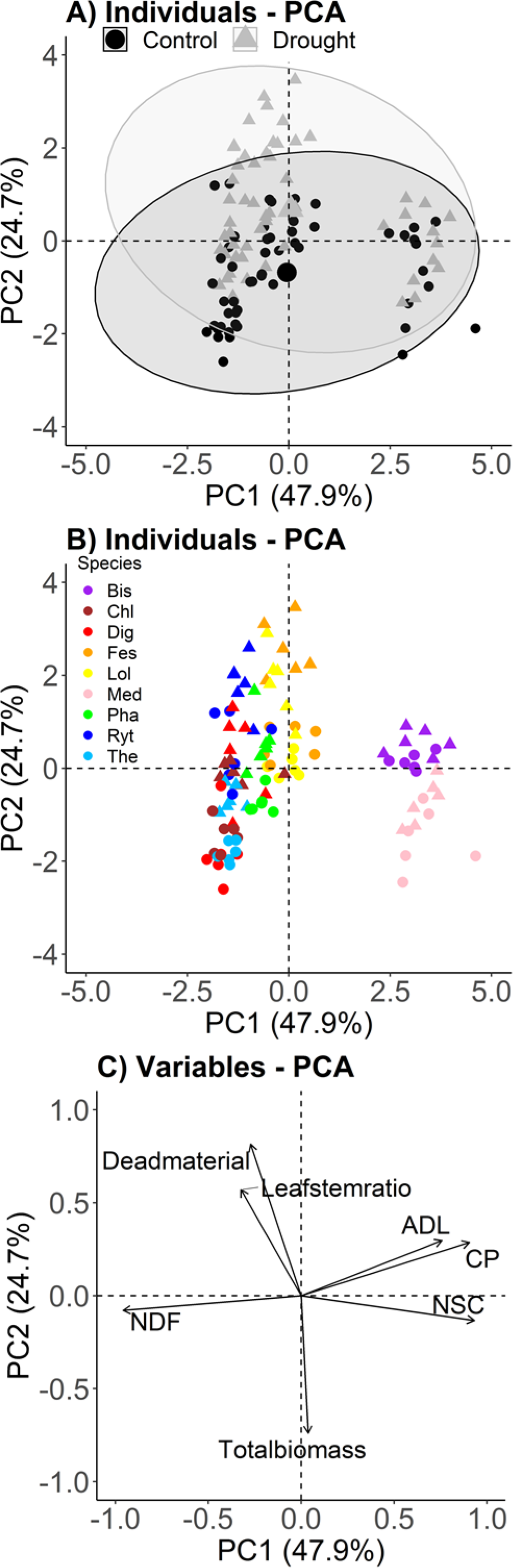
Principal component biplots illustrating variation in total biomass production, structural and nutritional traits across pasture species: A) scores for plant individuals grouped by treatment (control = circles and drought = triangles) with 95% confidence ellipses, B) scores for individuals, by species (colours; species abbreviations follow Figure 1); and C) variables loadings. Nutritional parameters abbreviations are as follows: crude protein (CP), non-structural carbohydrates (NSC), neutral detergent fibre (NDF), acid detergent lignin (ADL).

In general, the percentage of dead material and leaf:stem ratio were negatively associated with total biomass production, while CP, ADL and NSC were negatively associated with NDF (**Figure 3C**). The control treatment was associated with higher biomass and the drought treatment with more dead material and a high leaf:stem ratio. Furthermore, the percentage of dead material and leaf:stem ratio were higher for C3 grasses; high concentrations of ADL, CP and NSC were associated with legumes, while high values of NDF were associated with C4 grasses.

## 4 DISCUSSION

Here, we present the effects of a 6-month winter/spring drought on productivity as well as aboveground plant structure and nutritional composition for a diverse range of globally-important pasture species. Consistent with our hypothesis, we found that drought reduced pasture productivity and increased dead material across multiple harvests, although some species were unaffected during individual harvests. In most species, ongoing drought amplified the negative effects on productivity from one harvest to the next, but not on nutritional quality. There were large differences in the magnitude and direction of species’ responses to water stress in terms of leaf:stem biomass ratios and nutritional quality. In some cases, these findings were contrary to our expectations of reduced nutritional quality under drought. Significant changes to whole-plant nutritional quality were generally driven by simultaneous changes to both leaf and stem tissues, but in a few species were associated with changes in only one tissue. Across the entire experiment, *Chloris*, *Lolium* and *Rytidosperma* were the species most adversely impacted by drought in terms of productivity, dead material and nutritional quality, while *Biserrula* and *Themeda* were the least affected. *Medicago* and *Phalaris* were the only species with no change in nutritional quality in response to drought. The species-specific nature of morphological and nutritional responses to drought highlights the importance of carrying out studies across multiple plant species, with diverse traits, to better understand climate change impacts on pastures.

### 4.1 Productivity and dead material

Change in aboveground productivity is a fundamental plant response to environmental change (Wang et al., 2007). Studies have highlighted the impacts of drought on biomass reduction across pasture species, however, uncertainty remains in terms of the magnitude of the effects, and the consequences for production systems and/or ecosystem function (Cantarel., et al., 2013; Deleglise et al., 2015; Grant et al., 2014). Declining soil water content reduces plants’ ability to acquire sufficient water and nutrients for normal functioning, resulting in lower rates of plant growth and, in severe cases, causing tissue death (Buxton, 1996; Bruinenberg et al., 2002; Durand et al., 2010; Ren et al., 2016). The physiological mechanisms underpinning growth responses are often species-specific and reflect different strategies associated with drought resistance and drought survival (Baruch, 1994; Guenni et al., 2002; Munné-Bosch and Alegre, 2004).

Our observed reductions in productivity of up to 80% and increases in the percentage of dead material of up to 8-fold are aligned with previous studies exposing grassland species to short/long-term or moderate/severe drought conditions, which have reported large declines in biomass production (Cantarel et al., 2013; Deleglise et al., 2015) and increases in dead biomass (Power et al., 2016; Skinner et al., 2004). The great increase in the percentage of dead material in most of the species in this study may be due to both advanced senescence and a more rapid life cycle, as previously reported in severe drought stress scenarios (Bruinenberg et al., 2002; Ren et al., 2016). In our study, *Digitaria* and *Phalaris* showed consistent reductions in production and increases in dead material across the 6-month drought treatment, whereas other species had responses that differed between harvests. There are a number of mechanisms that might drive such differences, including different drought sensitivities at various stages in the plant’s life cycle or different degrees of realised water stress– reflecting the actual timing of rain (irrigation) events and temperature differences driving potential evapotranspiration, at different stages in winter and spring. In addition, these temporally variable effects of drought align with research emphasizing plant species’ adjustments in growth and resource allocation during exposure to drought conditions (Eziz et al., 2017; Gray and Brady, 2016). For example, some species may accumulate nutrients that were not used for growth during a drought event, but then are available for a rapid increase in leaf growth during any rewatering event that preceded a specific harvest, as reported by Guenni et al. (2002) in a study with forage grass species. Overall, our findings highlight species differences in ability to tolerate and adapt to drought, as well as seasonal/phenology effects on the extent of drought sensitivity (Gray and Brady, 2016; Lee et al., 2013).

### 4.2 Nutritional composition and structural biomass allocation

Reduced growth and increased senescence and/or death of biomass during drought have been reported to significantly affect the nutritional quality of forage species (Deleglise et al., 2015; Dumont et al., 2015; Ren et al., 2016). The proportion of dead material influences forage nutritional quality, as dead herbage is always associated with low forage energy value and digestibility (Hodgson et al., 1990; Shakhane et al., 2013). We found a significant negative correlation between the percentage of dead material and digestibility across all pasture species from both watering regimes throughout the experimental period (**Supplementary Figure S2A**). While lower forage nutritional quality and digestibility are often reported in response to severe drought (Buxton, 1996; Deleglise et al., 2015; Durand et al., 2010; Ren et al., 2016), no change or slight improvements in quality are commonly reported in response to moderate drought (Dumont et al., 2015; Kuchenmeister et al., 2013; Staniak and Harasim, 2018).

In this study, the drought-related decrease in nutritional quality associated with increased fibre – mainly the lignin fraction – may be explained by plant maturation, leaf senescence and cellular modifications that certain species develop to prevent water losses (Habermann et al., 2021; Le Gall et al., 2015). Previous studies have reported that under severe drought stress, as plant maturation accelerates, stem growth advances, thereby decreasing the leaf:stem ratio and increasing the accumulation of fibrous components, which may result in forage toughness and lower forage quality and digestibility (Bruinenberg et al., 2002; Deetz et al., 1996; Dumont et al., 2015; Ren et al., 2016). The hypothesis that the reduction in nutritional quality due to severe drought would be associated with a decrease in leaf:stem ratio was confirmed only for one species (*Chloris*) in our study. In addition, we did not find a correlation between leaf:stem ratio and digestibility under drought conditions among the pasture species throughout the experimental period, although there was a positive correlation under control conditions (**Supplementary Figure S2B**). Furthermore, increases in fibrous components in some plant species under severe drought, such as accumulation of lignin (an important component of the plant cell wall) can reduce plant cell wall water penetration and transpiration, helping to maintain cell osmotic balance and protect membrane integrity under drought stress (Liu et al., 2018b, Moura et al., 2010). However, this may have important implications for animal nutrition as lignin acts as a barrier to fibre degradation by rumen microbes, making energy from fibre unavailable for ruminants and ultimately decreasing forage digestibility (Amiri et al., 2012; Buxton et al., 1995; Grev et al., 2020; Jung et al., 1997).

In our study, although we found that under drought, *Chloris* and *Lolium* significantly increased lignin (up to +103% and +63%, respectively; November harvest), only *Lolium* decreased digestibility when compared to the control treatment (**Supplementary Figure S3**). However, *Lolium* digestibility was still within the digestibility range (60-70%) required for maintaining moderate livestock production (DPI, 2020). These findings indicate that drought-induced changes in nutritional composition may still result in some species being able to provide sufficient nutrients to maintain digestion process and moderate animal production. The only exception to this in our study was for *Chloris*, a C4 grass, in which CP (∼ 6.5%) was insufficient, even under control conditions, to ensure adequate fermentation and thus might reduce nutrient utilization efficiency by the ruminal microbiota and negatively affect animal production (NRC, 2001; Van Soest, 1994). If *Chloris* were to be used as pasture, it would need to be used in conjunction with high-protein food, such as legume species or urea supplementation, to optimize nutrient use efficiency and production goals (e.g. liveweight gains or milk production), even when grown under higher rainfall conditions.

Importantly, for a subset of our species, we found an increase in nutritional quality under drought through an increase in CP and NSC, and a decrease in NDF and ADL. Previous studies have reported that moderate drought stress can induce a delay in plant maturation and growth, resulting in plants with fewer, shorter stems and flowering parts, and increases in leaf:stem ratio, which explains much of the improved crude protein concentrations and digestibility (Buxton, 1996; Dumont et al., 2015; Kuchenmeister et al., 2013; Staniak and Harasim, 2018). In this context, the choice of species and varieties of pastures with delayed onset of flowering may allow for improved digestibility in drought conditions by increasing the leaf:stem ratios (Power et al., 2020). In our study, *Digitaria* and *Themeda* (both C4 grasses) increased allocation to leaves relative to stems under drought, and whole-plant digestibility subsequently increased when compared to the control treatment (November harvest; **Supplementary Figure S3)**. Furthermore, the increase in CP concentrations of some species under drought may be explained by trade-offs between nutrient accumulation and growth dilution, such that lower biomass production increased the tissue nitrogen concentration, as has also been reported in previous studies (Dumont et al., 2015; Grant et al., 2014).

In relation to the observed increases in non-structural carbohydrates, earlier studies with grasses suggest that this may alter the leaf osmotic potential, helping to maintain the uptake of soil water and thus resulting in increased drought tolerance and survival (DaCosta and Huang 2006; Fariaszewska et al., 2020; Volaire and Leliévre 1998). In our study, the reduced fibre and lignin concentrations found in some species (e.g. *Themeda)* can be explained by delayed stem elongation associated with slower rates of maturation and growth under water stress, as reported in previous studies (Buxton, 1996; Küchenmeister et al., 2013; Wilson et al., 1983). Such reduced stem elongation of some species under drought may result in higher leaf:stem biomass ratios, improving forage digestibility and sward structure for ease of grazing and forage intake (Buxton, 1996; Wilson et al., 1983).

In general, we found that while the direction and magnitude of drought impacts on forage nutritional quality varied across species and harvests, most of the pasture species were still able to provide nutrients to support ungulate digestion and, subsequently, maintain moderate animal production. However, a significant reduction in biomass production was common for all of the study species. This suggests that even with adequate forage nutritional quality, the amount of available forage may be insufficient to support the high performance of grazing ruminants in drought scenarios. In this case, reduced stocking densities may be an appropriate management strategy, although this would need species-level evaluation in future research studies.

## 5 CONCLUSIONS

The 6-month period of severe drought resulted in divergent responses in forage production, structural traits and nutritional composition among the nine pasture species examined. In general, productivity and percentage of dead material were more strongly and adversely impacted by drought than nutritional quality across all species. The changes in nutritional composition appeared to be related to either shifts in plant morphology (leaf:stem biomass ratios) or reduced growth, both of which were species-dependent, reflecting diverse drought adaptation strategies among species. Identification of the factors that drive changes in forage nutritional quality across different pasture species in response to various drought scenarios is essential to generating information about potential risks for farmers and industries in the face of climate change. This knowledge can inform management strategies in relation to the timing of grazing or cutting, selection of drought-tolerant species/cultivars, and optimization of forage resources to support animal performance. Future research is needed with animal trials to determine the extent to which observed changes in the nutritional quality of pasture species affect forage intake and animal production (e.g. milk and meat), as well as the incidental environmental impacts of consuming forage produced under drought conditions, such as altered ruminant methane emissions – a key industry consideration.

## ACKNOWLEDGEMENTS

We are grateful to all the team members in Pastures and Climate Extremes (PACE) project specially Chioma Igwenagu, Gil Won Kim, Manjunatha Chandregowda and Vinod Jacob, as well as the following WSU summer scholars: Alexandra Boyd, Minh Doan, Samantha Weller, Shania Therese Didier Serre and Ben Capel. This work was supported by funding from the Meat & Livestock Australia Donor Company (P.PSH.0793), Dairy Australia (C100002357) and Western Sydney University. The authors would like to thank the technical team (Burhan and Craig B) at Western Sydney University for technical support. The authors declare that they have no conflicts of interest regarding the publication of this article.

## AUTHOR CONTRIBUTIONS

KLMC, ACC, HZ and KJF performed the experiment. KLMC processed, analysed the samples, conducted statistical analyses (with input from ACC) and drafted the manuscript. All the co-authors designed the experiment and provided input on subsequent drafts.

## DATA AVAILABILITY

The data that support the findings of this study are available from the authors upon reasonable request.

