## APPENDIX for "Plant structural and nutritional responses to drought differ among common pasture species"

### SUPPLEMENTARY DATA

**Table S1.** Plant biomass harvests by pasture species performed during the drought period of 2019 (August, October, November); measurements taken (productivity, percentage of dead material, leaf:stem ratio) and plant materials (whole-plant, leaves and stems) used for nutritional analysis from each harvest.

| **Species** | **Months** | | |
|---|---|---|---|
| | **August** | **October** | **November** |
| *Biserrula pelecinus* | Harvest | Harvest | Harvest |
| *Chloris gayana* | NA | Harvest | Harvest |
| *Digitaria eriantha* | NA | Harvest | Harvest |
| *Festuca arundinacea* | Harvest | Harvest | Harvest |
| *Lolium perenne* | Harvest | Harvest | Harvest |
| *Medicago sativa* | Harvest | Harvest | Harvest |
| *Phalaris aquatic* | Harvest | Harvest | Harvest |
| *Rytidosperma caespitosum* | Harvest | Harvest | Harvest |
| *Themeda triandra* | Harvest | Harvest | Harvest |
| Measurements taken: | Productivity<br>Dead material | Productivity<br>Dead material | Productivity<br>Dead material<br>Leaf:stem ratio |
| Plant materials used for nutritional analysis: | Whole-plant | Whole-plant | Whole-plant<br>Leaves<br>Stems |

*NA: no harvest due to lack of plant biomass in the plots of these species.*

**Table S2.** *P* values for productivity, percentage of dead material, and nutritional composition of the whole-plant of pasture species in response to drought treatment during the drought period (August, October and November harvests).

| Species | Harvest | Productivity | Dead material | CP | NSC | NDF | ADL |
|---|---|---|---|---|---|---|---|
| Bis | August | **0.04** | 0.29 | 0.59 | 0.91 | 0.95 | 0.47 |
| | October | **0.02** | **0.04** | **0.05** | 0.15 | 0.43 | 0.73 |
| | November | 0.38 | 0.49 | **<0.01** | 0.17 | 0.61 | 0.35 |
| Chl | October | 0.05 | 0.52 | 0.68 | 0.68 | 0.56 | 0.61 |
| | November | **<0.01** | **<0.01** | 0.31 | 0.06 | 0.50 | **<0.01** |
| Dig | October | **<0.01** | **<0.01** | **0.05** | 0.81 | **0.05** | 0.67 |
| | November | **<0.01** | **0.02** | **<0.01** | 0.43 | **<0.01** | 0.27 |
| Fes | August | 0.67 | 0.69 | **<0.01** | 0.51 | 0.15 | 0.06 |
| | October | **<0.01** | **<0.01** | 0.08 | 0.05 | 0.50 | 0.22 |
| | November | 0.63 | **<0.01** | **<0.01** | 0.59 | 0.28 | **<0.01** |
| Lol | August | 0.09 | 0.75 | **0.03** | 0.18 | 0.72 | 1.00 |
| | October | **<0.01** | **0.03** | **<0.01** | **<0.01** | 0.55 | 0.14 |
| | November | **<0.01** | **<0.01** | 0.19 | 0.13 | **<0.01** | **<0.01** |
| Med | August | 0.78 | 0.75 | 0.93 | 0.18 | 0.13 | 0.50 |
| | October | **<0.01** | 0.97 | 0.13 | 0.14 | 0.93 | 0.65 |
| | November | **<0.01** | **0.04** | 0.12 | 0.09 | 0.78 | 0.86 |
| Pha | August | **0.03** | **0.01** | 0.97 | 0.40 | 0.51 | 0.20 |
| | October | **<0.01** | **0.01** | 0.16 | 0.06 | 0.54 | 0.39 |
| | November | **<0.01** | **<0.01** | 0.07 | 0.45 | 0.85 | 0.08 |
| Ryt | August | 0.10 | 0.32 | 0.47 | 0.22 | 0.57 | 0.89 |
| | October | **<0.01** | **<0.01** | 0.06 | 0.59 | **0.05** | **0.01** |

| | | | | | | | |
|---|---|---|---|---|---|---|---|
| | November | 0.15 | **<0.01** | 0.70 | 0.55 | 0.12 | **0.04** |
| The | August | 0.08 | 0.44 | 0.45 | 0.50 | 0.43 | 0.74 |
| | October | **<0.01** | 0.95 | 0.25 | **<0.01** | 0.06 | 0.28 |
| | November | **<0.01** | 0.81 | 0.72 | **0.04** | 0.06 | **0.02** |

*Note:* **Bold** *values denote statistical significance at the p ≤ 0.05 level. During the harvest in August, there was no biomass for Chloris and Digitaria.*

*Abbreviations: CP: crude protein; NSC: non-structural carbohydrates; NDF: neutral detergent fibre; ADL: acid detergent lignin; Bis: Biserrula; Chl: Chloris; Dig: Digitaria; Fes: Festuca; Lol: Lolium; Med: Medicago; Pha: Phalaris; Ryt: Rytidosperma; The: Themeda.*

**Table S3.** *P* values for leaf:stem ratio and nutritional composition of plant parts (leaves and stems) of pasture species in response to drought treatment at the end of the drought period (November harvest).

| Variables | Species | | | | | | | | |
|---|---|---|---|---|---|---|---|---|---|
| | ***Bis*** | ***Chl*** | ***Dig*** | ***Fes*** | ***Lol*** | ***Med*** | ***Pha*** | ***Ryt*** | ***The*** |
| Leaf:stem ratio | 0.47 | **0.01** | **0.03** | 0.44 | 0.67 | 0.84 | **<0.01** | 0.35 | **<0.01** |
| CP | | | | | | | | | |
| Leaves | **<0.01** | 0.11 | **<0.01** | **0.02** | 0.35 | 0.06 | 0.64 | 0.96 | 0.10 |
| Stems | 0.08 | 0.25 | **<0.01** | 0.35 | **0.04** | 0.84 | 0.35 | 0.34 | 0.96 |
| NSC | | | | | | | | | |
| Leaves | 0.06 | 0.08 | 0.30 | 0.43 | 0.21 | 0.08 | **0.02** | 0.37 | **0.05** |
| Stems | 0.09 | 0.48 | 0.41 | 0.76 | **0.01** | **0.03** | **<0.01** | 0.17 | **0.05** |
| NDF | | | | | | | | | |
| Leaves | 0.60 | 0.99 | **<0.01** | 0.63 | **<0.01** | 0.98 | **<0.01** | 0.56 | 0.62 |
| Stems | 0.39 | 0.97 | **<0.01** | **0.05** | 0.08 | 0.07 | **<0.01** | 0.93 | 0.12 |
| ADL | | | | | | | | | |
| Leaves | 0.69 | **<0.01** | 0.28 | **0.04** | **<0.01** | 0.79 | 0.47 | 0.98 | 0.30 |
| Stems | 0.41 | **<0.01** | 0.13 | **0.01** | **<0.01** | 0.44 | 0.34 | **<0.01** | **0.05** |

*Note:* ***Bold*** *values denote statistical significance at the $p \leq 0.05$ level.*

*Abbreviations follow Table S2.*

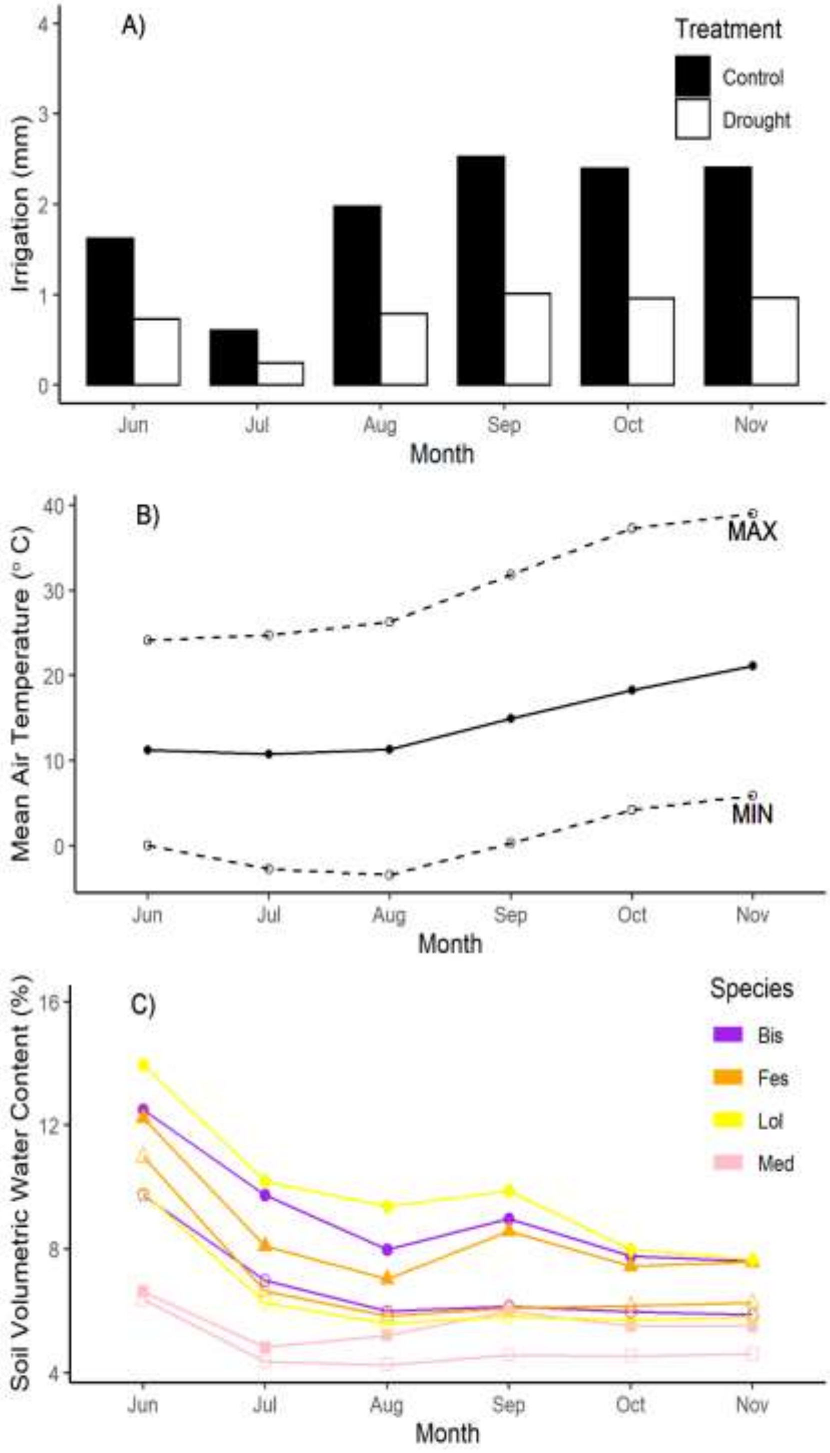


**Figure S1.** Amount of irrigation (mm; panel A) applied in each treatment (control, drought), mean air temperature (maximum and minimum; panel B) and soil moisture content in four species subplots (*Biserrula*, *Festuca*, *Lolium*, *Medicago*; panel C) under both treatments (control: filled shape and drought: empty shape) during the experimental period (1 June to 30 November 2019).

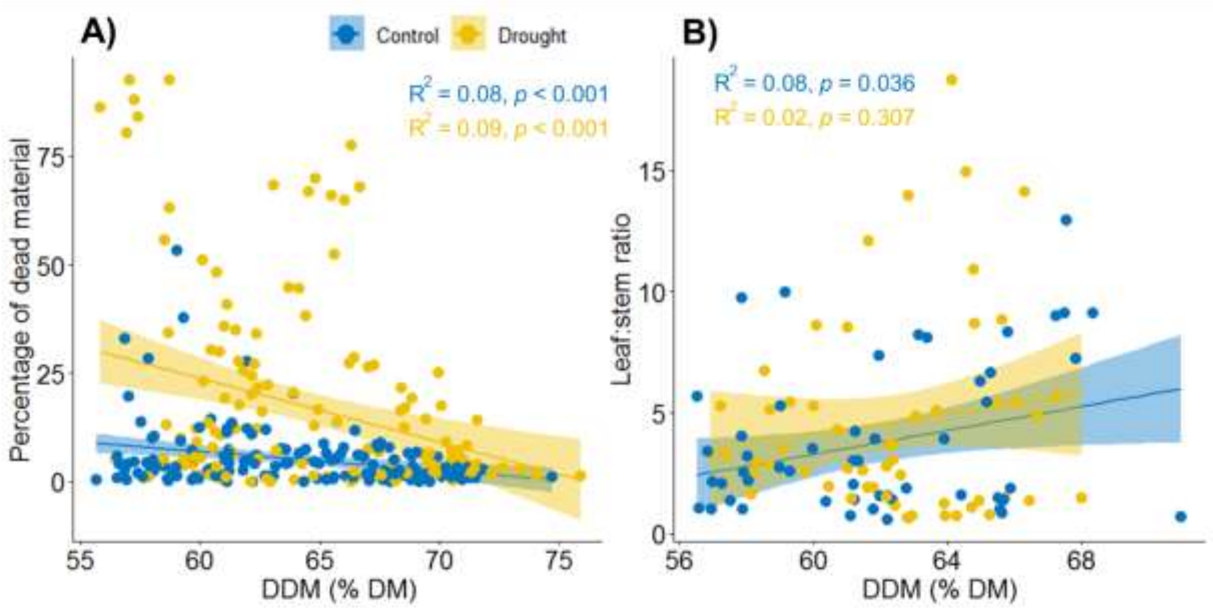


**Figure S2.** Correlations between A) percentage of dead material, B) leaf:stem ratio and estimated digestible dry matter (DDM in % of dry matter; whole-plant) for the nine pasture species studied under different watering regime treatments (control and drought) during the 6-month experimental period (August, October and November harvests). Correlations were tested with Pearson correlation, with $R^2$ and $p$ values for each treatment shown in different colors in each panel (blue = control, yellow = drought). DDM was calculated according to Oddy et al. (1983) as follows: DDM % = 83.58 - 0.824 ADF% + 2.626 N%.

*Reference: Oddy, V.H., Robards G.E., Low, S.G., 1983. Prediction of in vivo digestibility of ruminant feeds from fibre and nitrogen content, in: Robards, G.E., Packham, R.G. (Eds.), Feed information and animal production: Proceedings of the 2nd international conference of international network of feeds information centres, Farnham Royal, UK: Commonwealth Agricultural Bureaux, pp. 395-398.*

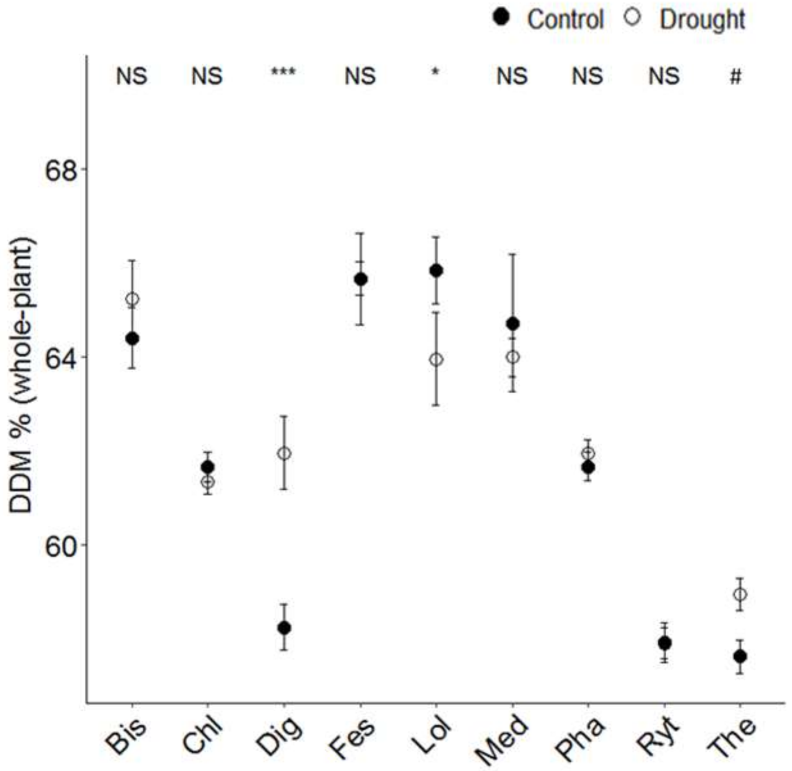


**Figure S3.** Estimated digestible dry matter (DDM %; whole-plant) for the nine pasture species studied under different watering regimes treatments (closed circle = control, open circle = drought) in November harvest. DDM calculation follows Figure S2. Significant comparisons for the effects of treatments are indicated as follows: NS= not significant, # $p \leq 0.1$, * $p < 0.05$, *** $p < 0.001$. Species abbreviations are as follows: *Biserrula* (Bis), *Chloris* (Chl), *Digitaria* (Dig), *Festuca* (Fes), *Lolium* (Lol), *Medicago* (Med), *Phalaris* (Pha), *Rytidosperma* (Ryt), *Themeda* (The).
